# Aryl hydrocarbon receptor (AhR) activation by 2,3,7,8-tetrachlorodibenzo-*p*-dioxin (TCDD) dose-dependently shifts the gut microbiome consistent with the progression of steatosis to steatohepatitis with fibrosis

**DOI:** 10.1101/2021.11.02.466980

**Authors:** Russell R. Fling, Tim R. Zacharewski

## Abstract

Gut dysbiosis with disrupted enterohepatic bile acid metabolism is commonly associated with non-alcoholic fatty liver disease (NAFLD) and recapitulated in a NAFLD-phenotype elicited by 2,3,7,8-tetrachlorodibenzo-p-dioxin (TCDD) in mice. TCDD induces hepatic fat accumulation and increases levels of secondary bile acids including taurolithocholic acid and deoxycholic acid, microbial modified bile acids involved in host bile acid regulation signaling pathways. To investigate the effects of TCDD on the gut microbiota, cecum contents of male C57BL/6 mice orally gavaged with sesame oil vehicle or 0.3, 3, or 30 µg/kg TCDD were examined using shotgun metagenomic sequencing. Taxonomic analysis identified dose-dependent increases in Lactobacillus species (i.e., *Lactobacillus reuteri*). Increased species were also associated with dose-dependent increases in bile salt hydrolase sequences, responsible for deconjugation reactions in secondary bile acid metabolism. Increased *L. reuteri* levels were further associated with mevalonate-dependent isopentenyl diphosphate (IPP) biosynthesis and menaquinone biosynthesis genes. Analysis of gut microbiomes from cirrhosis patients identified increased abundance of these pathways as identified in the mouse cecum metagenomic analysis. These results extend the association of lactobacilli with the AhR/intestinal axis in NAFLD progression and highlight the similarities between TCDD-elicited phenotypes in mice to human NAFLD.

## 1. Introduction

Non-alcoholic fatty liver disease (NAFLD) is estimated to affect ∼25% of the global population and is defined as a spectrum of progressive pathologies that include steatosis, immune cell infiltration/inflammation, fibrosis, and cirrhosis. It is associated with increased risk for hepatocellular carcinoma, and is the 2^nd^ leading cause of liver transplants in the United States[1]. Other pathologies including obesity, type 2 diabetes (T2D), and coronary heart disease demonstrate high co-occurrence with NAFLD, e.g., ∼40-70% in T2D patients and ∼90% in obese patients[2]. A multi-hit hypothesis for NAFLD proposes several contributing factors to development and progression including disruptions in the immune system, adipose tissue metabolism, and the gut microbiome[3]. Emerging evidence also suggests environmental contaminants may play an underappreciated role in gut dysbiosis and NAFLD development[4–11]. Specifically, 2,3,7,8-tetrachlorodibenzo-*p*-dioxin (TCDD), a persistent environmental organochloride pollutant, induces steatosis and the progression to steatohepatitis with fibrosis in mice resembling human NAFLD development[9,12–14]. TCDD-induced dyslipidemia also exhibits other similar NAFLD characteristics such as decreased VLDL secretion, free fatty acid accumulation, inhibition of β-oxidation, and disrupted cholesterol and bile acid metabolism[9,15–18].

The effects of TCDD and other related polychlorinated dibenzo-p-dioxins (PCDDs), dibenzofurans (PCDFs) and biphenyl (PCBs) as well as polyaromatic hydrocarbons (PAHs), are mediated through activation of the aryl hydrocarbon receptor (AhR), a basic helix-loop-helix/Per-Arnt-Sim transcription factor typically associated with xenobiotic metabolism[19]. In addition, the AhR plays an essential role in gut homeostasis through regulation of the immune system and bile acid metabolism[9,15,20,21], with endogenous and xenobiotic AhR ligands affecting the gut microbiome congruent with NAFLD-like pathology[8–10]. Moreover, gut dysbiosis is commonly reported in NAFLD, making the gut microbiome an attractive target for non-invasive diagnostic tools and potential target for intervention[22,23].

Although the AhR exhibits promiscuous binding activity for a wide variety of structurally diverse xenobiotics, natural products, and endogenous metabolites, its endogenous role remains unknown[24]. Upon ligand binding, the cytosolic AhR disassociates from chaperone proteins and translocates to the nucleus where it dimerizes with the AhR nuclear transporter (ARNT). The AhR/ARNT heterodimer complex then binds to dioxin response elements located throughout the genome, affecting gene expression[25]. Endobiotic ligands for the AhR include host-derived metabolites such as tryptophan catabolites (e.g., L-kynurenine), microbial-produced indole derivatives (e.g., indole-3-aldehyde produced by *Lactobacillus reuteri*), and compounds derived from fruits and cruciferous vegetables (e.g., 6-formylindolo[3,2-b]carbazole [FICZ])[24]. Microbial produced indoles activate AhR in the intestine, affecting barrier function and homeostasis by regulating the intestinal immune system through CD4^+^ T-cell differentiation, and the induction of interleukin (IL)-22 and IL-10 cytokine production[25]. AhR-dependent IL-22 induction subsequently increases antimicrobial peptides expression in intestinal epithelial cells, inhibiting pathogen infection and inflammation[25–27].

Knockout models and/or treatment with endogenous and xenobiotic AhR ligands results in shifts in the gut microbiome with diverse effects depending on the model and ligand[8,11,27–29]. Shifts in Firmicutes/Bacteroidetes ratio can differ between AhR ligands, e.g., 2,3,7,8-tetrachloro dibenzofuran decreased the ratio [8] whereas TCDD increased it[11]. However, responses in various AhR models are in agreement regarding increased secondary bile acids [8,9] and effects on segmented filamentous bacteria [8,11,28]. AhR knockout models, and treatment with TCDD or other endogenous compounds also demonstrate strong correlations between AhR activation and enrichment of Lactobacillus species, i.e., *L. reuteri* [27–31]. Tryptophan catabolism to AhR ligands by Lactobacillus species is a proposed mechanism for gut microbial regulation of AhR signaling that modulates intestinal and gut microbiome homeostasis[27].

Bile acids also affect the gut microbiome by exerting antimicrobial activity[32]. Conversely, the gut microbiota play critical roles in host bile acid homeostasis through microbial metabolism that qualitatively and quantitatively impact bile acid composition with consequences for bile acid activated signaling pathways in the host. The gut microbiome performs the first step of bile acid deconjugation with subsequent oxidation, reduction or dehydroxylation reactions to produce diverse secondary bile acid molecular species[33]. Select secondary bile acids ,*e.g*., glycodeoxycholic acid [GDCA], demonstrate higher inhibition of bacterial growth compared to other primary and secondary bile acids[34]. In regards to the host, some secondary bile acids ,*e.g*., lithocholic acid [LCA] and deoxycholic acid [DCA]), exhibit high affinity for the farnesoid x receptor (FXR) and G protein-coupled bile acid receptor (TGR5, *a.k.a*., GPBAR1), which regulate glucose, lipid, and bile acid homeostasis[35–37]. In human NAFLD, secondary bile acid metabolism is disrupted with bile acid analogs that target FXR and TGR5 signaling pathways under development for the treatment of liver disease [23,32,38].

Previous work has demonstrated serum levels of LCA and DCA increased following TCDD treatment suggesting enrichment for microbial bile acid metabolism[9]. To further explore dose-dependent disruptions in gut microbiome and microbial metabolism relevant to the progression of NAFLD-like pathologies, shotgun metagenomic analysis was used to examine the dose dependent taxonomic and metabolic disruptions elicited by TCDD.

## 2. Results

### 2. 1. TCDD-elicited toxicity enriched for Lactobacillus species

Taxonomic analysis identified significant dose-dependent population shifts among caecum microbiota in response to TCDD. While no significance was observed between treatment groups at the phylum level, a decreasing trend was observed for Bacteroidetes concurrent with increasing trends in Firmicutes abundance (Figure 1a).

**Figure 1.**
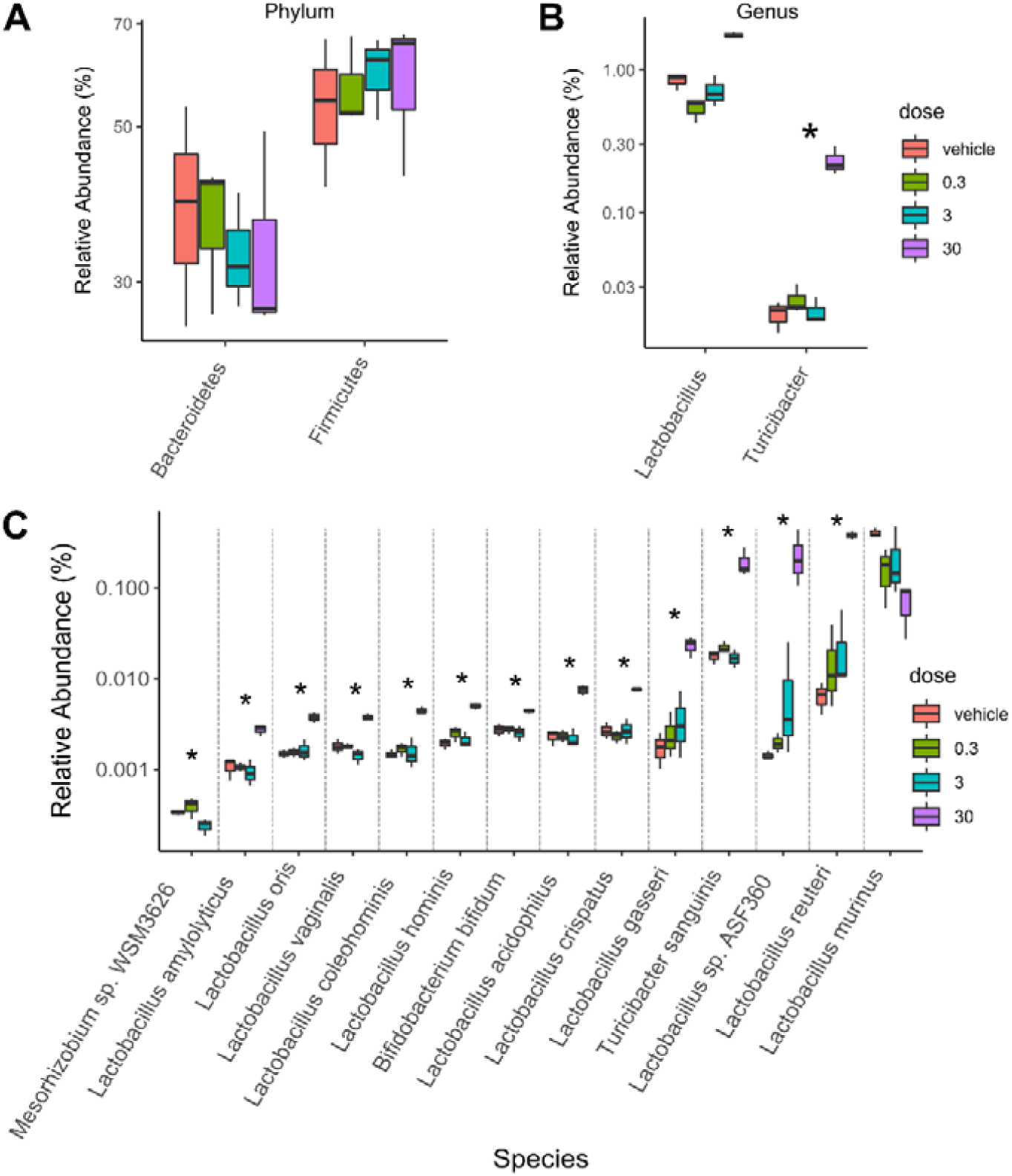
TCDD enriched Lactobacillus species in the cecum microbiota. Taxa abundance were assessed in metagenomic cecum samples from male C57BL/6 mice following oral gavage with sesame oil vehicle or 0.3, 3 or 30 µg/kg TCDD every 4 days for 28 days (n=3). Significant shifts in relative abundances of taxa are presented at the **(A)** phylum, **(B)** genus, **(C)** and species levels. Significance is denoted with an asterisk (*; adjusted p-value < 0.1).

At the genus level, Turicibacter was enriched by TCDD while the genus Lactobacillus trended towards enrichment (Figure 1b). Interestingly, at the species level, 10 out of 13 enriched species were from the Lactobacillus genus (*e.g*., *L. reuteri* and Lactobacillus sp. ASF360) as well as *Turicibacter sanguinis*. Conversely, the most abundant Lactobacillus species in vehicle treated mice, *Lactobacillus murinus*, trended towards a dose-dependent decrease (Figure 1c).

### 2.2 Bile salt hydrolase (*bsh*) levels correlated with significantly enriched species

Many Lactobacillus species deconjugate primary conjugated bile acids mediated by bile salt hydrolases (*BSH*), imparting bile acid tolerance[39]. To further investigate the effect of TCDD on bile acid metabolism, *bsh* sequences were annotated and quan ified within metagenomic samples. Annotations to *bsh* were increased by TCDD and associated with enriched species including *L. reuteri* and *T. sanguinis* (Figure 1c, Figure 2a, and Table S1).

**Figure 2.**
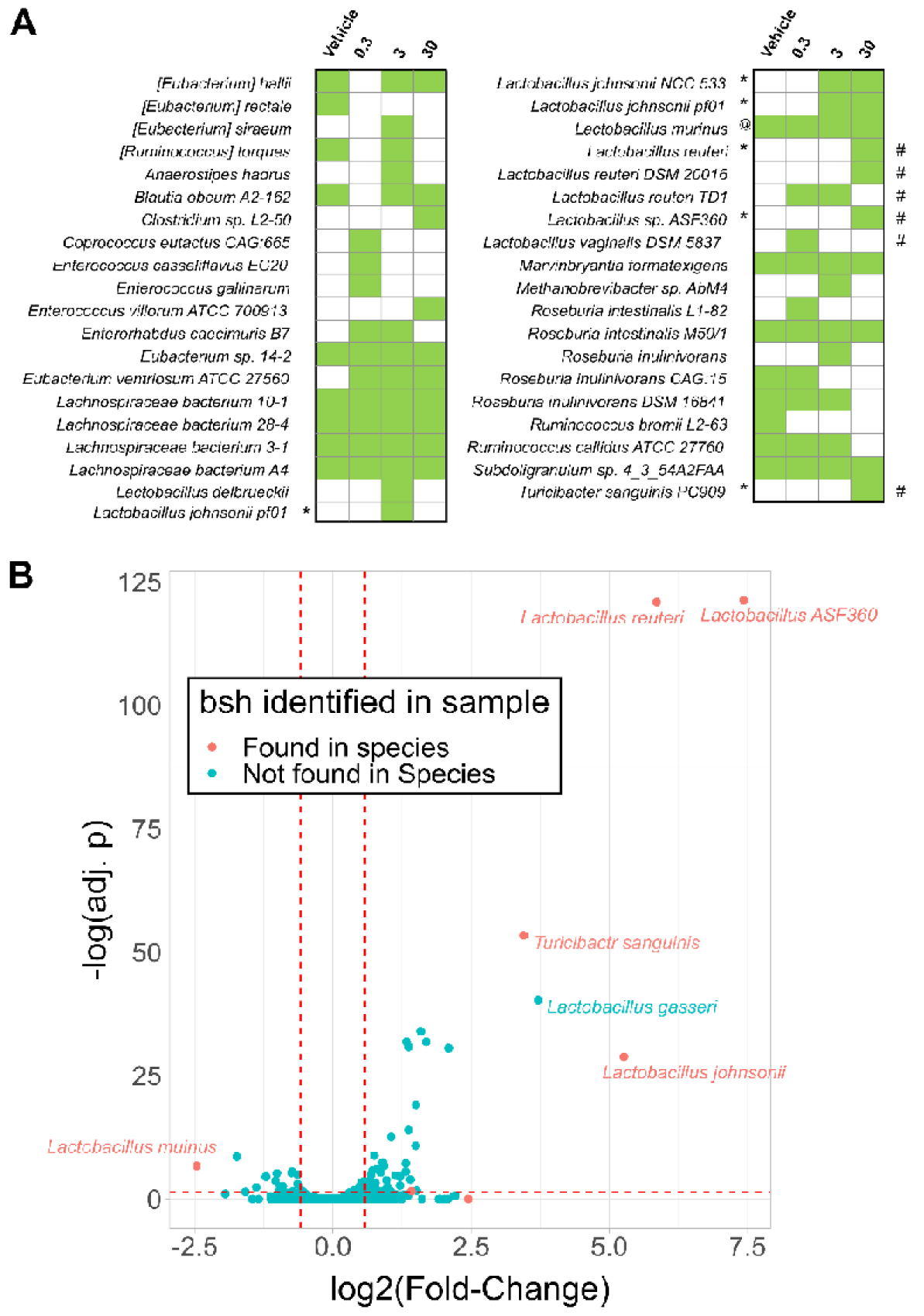
TCDD enriched Lactobacillus species possessing bile salt hydrolase (*bsh*). The presence of *bsh* gene sequences were assessed in metagenomic caecum samples from male C57BL/6 mice following oral gavage with sesame oil vehicle or 0.3, 3, or 30 µg/kg TCDD every 4 days for 28 days using 3 independent cohorts (n=3). **(A)** The presence (green boxes) or absence of *bsh* sequences detected in any of the metagenomic samples (n=3) are denoted within the respective treatment groups. Significant increases (*) or decreases (@) in normalized *bsh* abundances (adj. p < 0.1) are denoted. Also denoted is significantly increased species (#) determined by taxonomic analysis that corresponded with respective RefSeq species *bsh* annotations. Significance was determined by Maaslin2 R package. **(B)** Volcano plot displaying log2(fold-changes) in relative abundance of species between vehicle and 30 µg/kg TCDD treatment groups versus -log(adjusted p-values [adj. p]). Red dots denotes *bsh* sequences detected in 30 µg/kg TCDD treatment group. Significance was determined by the DeSeq2 R package comparing only vehicle and 30 µg/kg TCDD groups. Red dashed lines are reference to -log(0.05) value for y-axis and -1 and 1 for x-axis.

Conversely, *L. murinus* associated *bsh* annotations exhibited a dose-dependent decrease consistent with decreasing trends in taxonomic abundance. Although not reaching significance, many *bsh* sequences were also associated with unclassified Lachnospiraceae species including *Lachnospiraceae bacterium* A4, a community member reaching 5-23% relative abundance in the cecum metagenomic samples (Figure 2a). In contrast, *Lactobacillus gasseri* was enriched but no *bsh* sequences were identified (Figure 2b). To su marize, the top enriched species were also associated with increased abundances in *bsh* levels in the cecum.

### 2.3 TCDD enriched for mevalonate-dependent isoprenoid biosynthesis

To investigate other metabolic pathways imparting competitive advantages to TCDD-elicited gut environmental stresses, functional gene annotations associated with *L. reuteri*, the highest enriched species, were assessed. Among enriched uniref90 annotations in the cecum metagenomic dataset was the aromatic amino acid aminotransferase (UniRef90_A0A2S1ENB9) also classified to *L. reuteri* (Table S2). Aromatic amino acid aminotransferase produces a tryptophan metabolite, indole-3-aldehyde, a known AhR ligand reported to induce IL-22 *in vivo*[27]. Among 39 enzyme commission (EC) annotations that were enriched and associated with *L. reuteri* were several annotated to the isoprenoid biosynthesis pathway (Figure 3, Figure S1, and Table S3).

**Figure 3.**
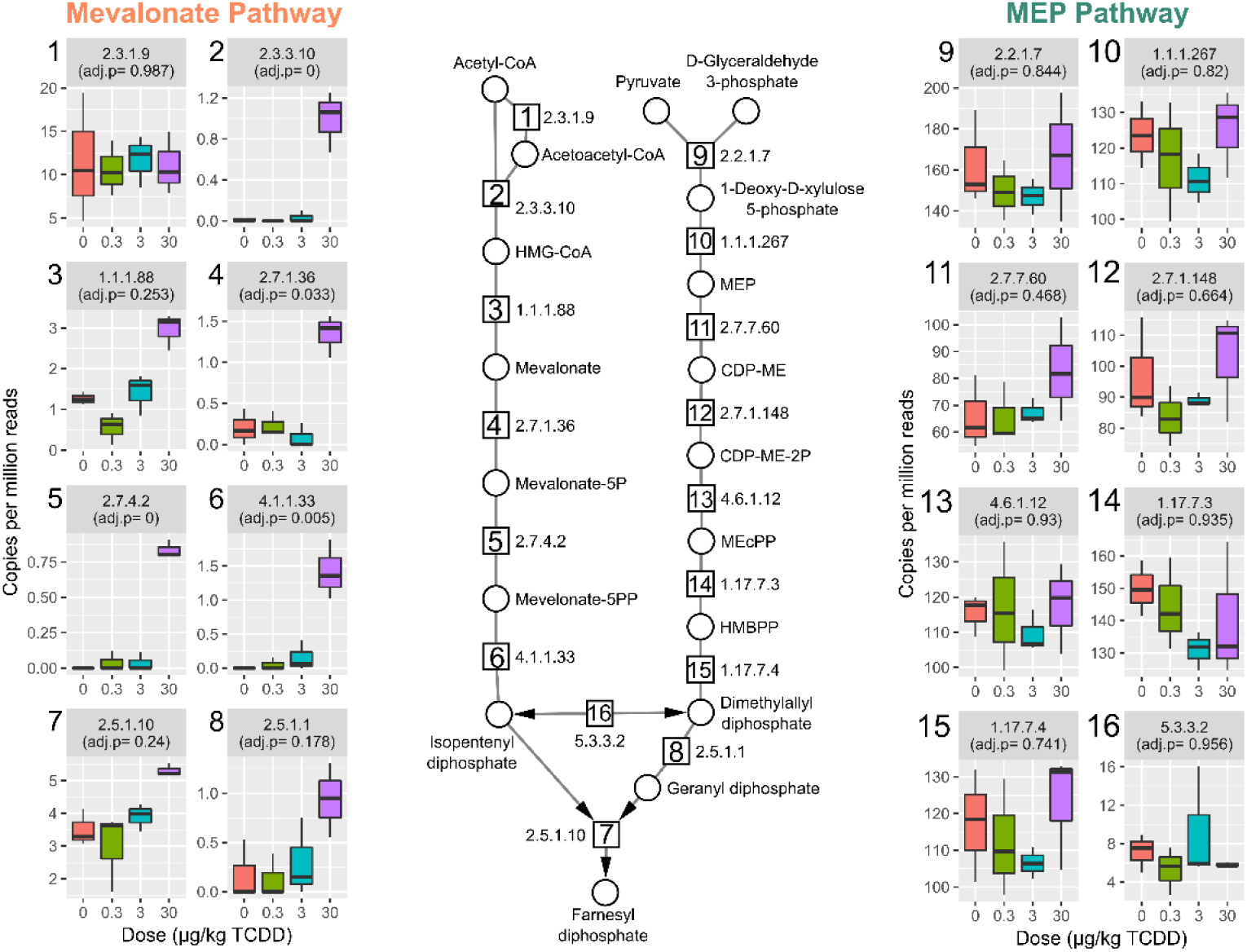
TCDD enriched genes from the mevalonate-dependent isoprenoid biosynthesis pathway. (a) Relative abundance of genes involved in isoprenoid biosynthesis and grouped by enzyme commission (EC) numbers for the mevalonate dependent and 2-C-methyl-D-erythritol 4-phosphate (MEP) pathways in cecum samples from male C57BL6 mice following oral gavage with sesame oil vehicle or 0.3, 3, or 30 µg/kg TCDD every 4 days for 28 days (n=3). Individual box plots are also numbered with the EC number matching the enzymatic step in pathway schematic. Adjusted p-values (adj. p) were determined by the Maaslin2 R package. Abbreviations.: 3-hydroxyl-3-methyl-clutaryl-CoA (HMG-CoA), (R)-5-Phosphomevalonate (mevalonate-5P), (R)-5-Diphosphomevalonate (mevalonate-5PP), 2-C-Methyl-D-erythritol 4-phosphate (MEP), 4-(Cytidine 5’-diphospho)-2-C-methyl-D-erythritol (CDP-ME), 4-(Cytidine 5’-diphospho)-2-C-methyl-D-erythritol (DEP-ME-2P), 2-C-Methyl-D-erythritol 2,4-cyclodiphosphate (MEcPP), 1-Hydroxy-2-methyl-2-butenyl 4-diphosphate (HMBPP).

Bacteria biosynthesize the isoprenoid, isopentenyl diphosphate (IPP), either through the mevalonate-dependent pathway, which is also found in mammals, or the methylerythritol phosphate (MEP)-pathway. Both *L. reuteri* and *Lactobacillus johnsonii* were the major contributors to mevalonate-dependent IPP biosynthesis pathway enrichment with almost all genes in the pathway increased by TCDD (Figure 3 and Figure S1). Gene enrichment in the alternative MEP-pathway were unchanged by TCDD. For *L. murinus*, only two EC annotations (EC 2.7.1.148, 4-Diphosphocytidyl-2-C-methyl-D-erythritol (CDP-ME) kinase, and EC 5.3.3.2, isopentenyl-diphosphate Delta-isomerase) were identified in the MEP pathway also found in *L. reuteri* (Figure S2).

HUMAnN 3.0 analysis of a published metagenomics dataset of fecal samples from human cirrhotic patients (https://www.ebi.ac.uk/ena/data/view/PRJEB6337) [40] revealed strikingly similar results to our caecum samples from TCDD treated mice. Specifically, increased gene abundance associated with the mevalonate-dependent pathways was also evident in patients with compensated and decompensated liver disease (Figure 4).

**Figure 4.**
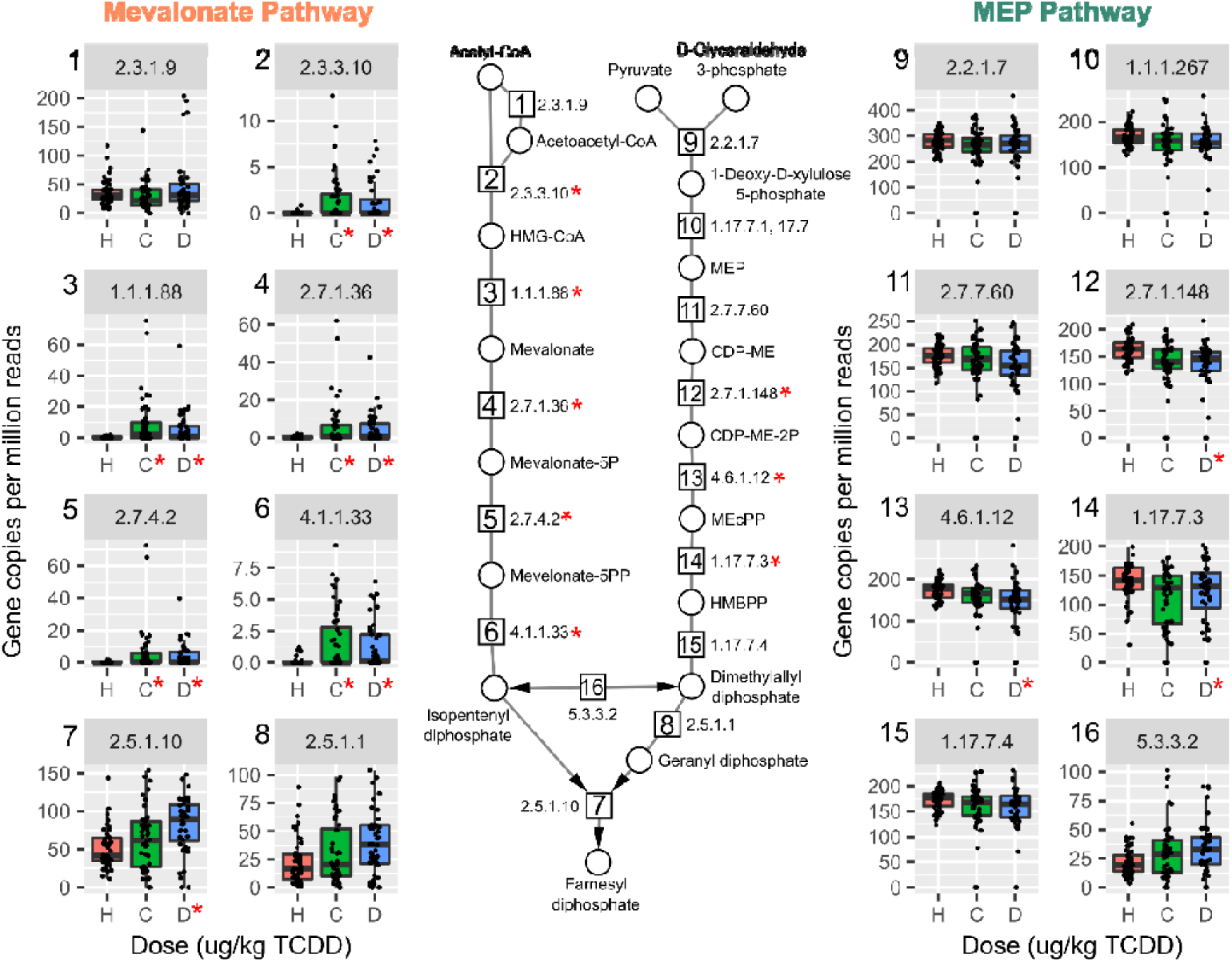
Mevalonate-dependent isoprenoid biosynthesis genes are enriched in a published metagenomics dataset of fecal samples from cirrhosis patients. Humann3 analysis of fecal gut microbiomes in healthy (H, red, n=52), compensated (C, green, n=48), or decompensated (D, blue, n=44) cirrhosis for mevalonate-dependent and methyl-D-erythritol 4-phosphate (MEP) pathways. Individual boxplots are numbered with the EC number matching the enzymatic step in pathway schematic. Significance is denoted with a red asterisk (*, adjusted p-values < 0.05) compared to healthy group. Abbreviations.: 3-hydroxyl-3-methyl-clutaryl-CoA (HMG-CoA), (R)-5-Phosphomevalonate (mevalonate-5P), (R)-5-Diphosphomevalonate (mevalonate-5PP), 2-C-Methyl-D-erythritol 4-phosphate (MEP), 4-(Cytidine 5’-diphospho)-2-C-methyl-D-erythritol (CDP-ME), 4-(Cytidine 5’-diphospho)-2-C-methyl-D-erythritol (DEP-ME-2P), 2-C-Methyl-D-erythritol 2,4-cyclodiphosphate (MEcPP), 1-Hydroxy-2-methyl-2-butenyl 4-diphosphate (HMBPP).

Compensated cirrhosis is defined as no decrease in liver function while decompensated cirrhosis exhibit decreased liver function. Among decompensated patients with cirrhosis, the mevalonate dependent IPP pathway was increased in 7 out of 8 EC numbers required for *de novo* IPP biosynthesis (Figure 4). Taxa annotated to genes in the pathway exhibited a wide variety in genera for each EC number in human samples compared to murine cecum samples from this study (Figure S3). Taxonomy classified to a majority of the mevalonate-dependent genes were from the Lactobacillaceae family including Enterococcus, Lactobacillus, Streptococcus genera (Figure S3 and Table S5). Lactobacillus and Streptococcus species including *L. reuteri* and *Streptococcus anginosus*, a known pathogen in liver abscesses[41], were among species classified to the pathway (Table S4 and Table S5).

### 2.4 Vitamin K2 (menaquinone) and peptidoglycan biosynthesis pathways in mouse NAFLD-phenotypes and gut microbiomes of cirrhosis patients

In polyprenol diphosphate biosynthesis, IPP is recursively added to geranyl diphosphate (GPP) or farnesyl diphosphate (FPP) for polyprenol biosynthesis used in vitamin K2 (*a.k.a*., menaquinone) and peptidoglycan biosynthesis [42,43]. TCDD enriched for heptaprenyl diphosphate synthase (EC 2.5.1.30) with major contributions from *L. reuteri* and *L. johnsonii* (Figure 5).

**Figure 5.**
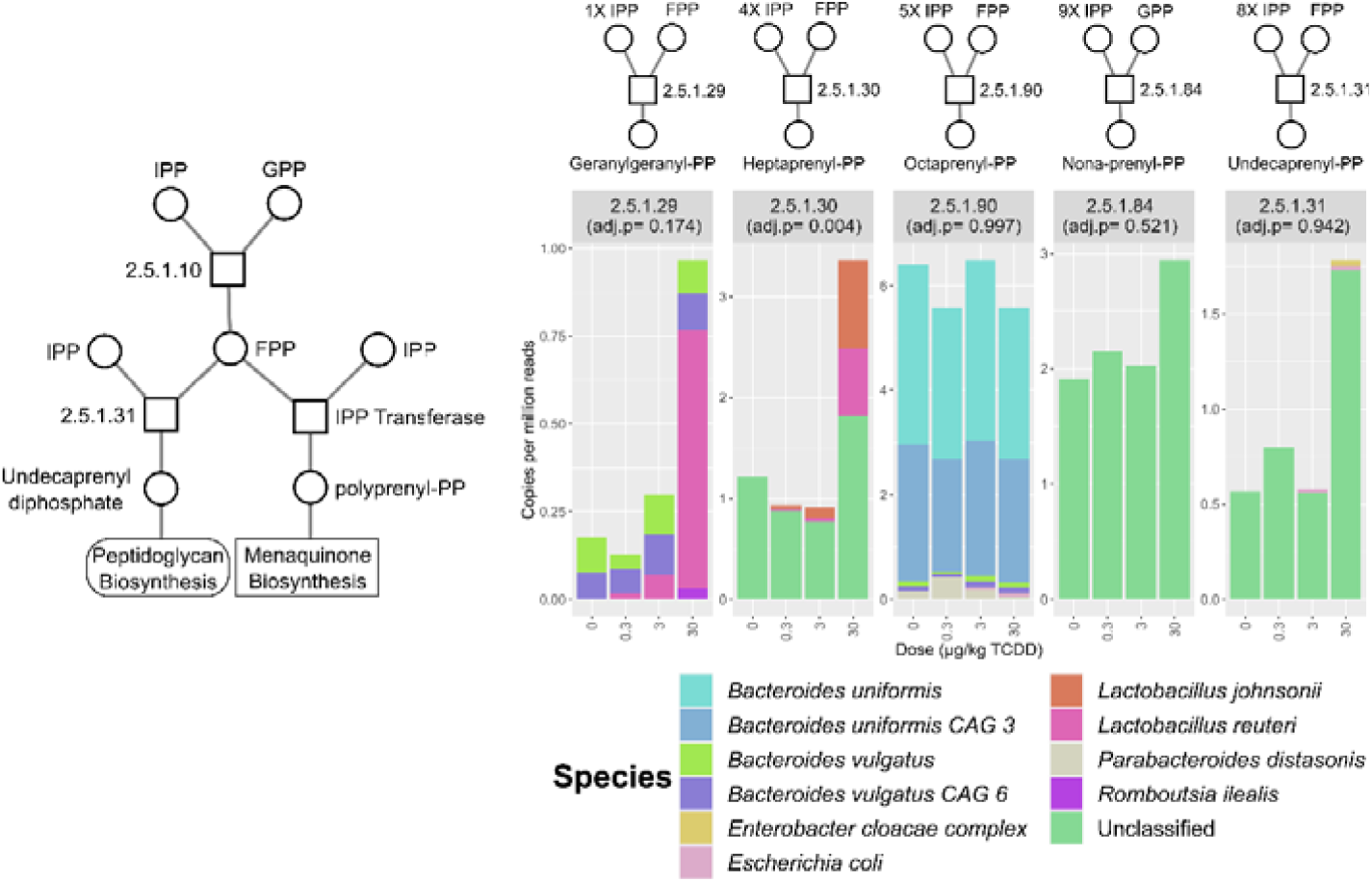
Relative abundance of polyprenol transferase EC annotations identified in the mouse cecum metagenomic dataset. Stacked bar plots represent mean relative abundance of grouped EC numbers (n=3) and represent identified species that contributed to mean total abundance for each treatment group. The number of isopentenyl diphosphate (IPP) and farnesyl diphosphate (FPP) molecules used for respective polyprenol biosynthesis are also denoted. Adjusted p-values were determined by the Maaslin2 R package. Abbreviations: isopentenyl diphosphate (IPP), geranyl diphosphate (GPP), polyprenyl diphosphate (polyprenyl-PP).

Because bacterial cell wall restructuring has been reported in response to bile acids and different levels of isoprenoid biosynthesis pathways were identified, peptidoglycan biosynthesis was also assessed[44]. Most genes encoding enzymes required for peptidoglycan biosynthesis were present in the metagenomic dataset (Figure 6a) with no changes observed following TCDD treatment.

**Figure 6.**
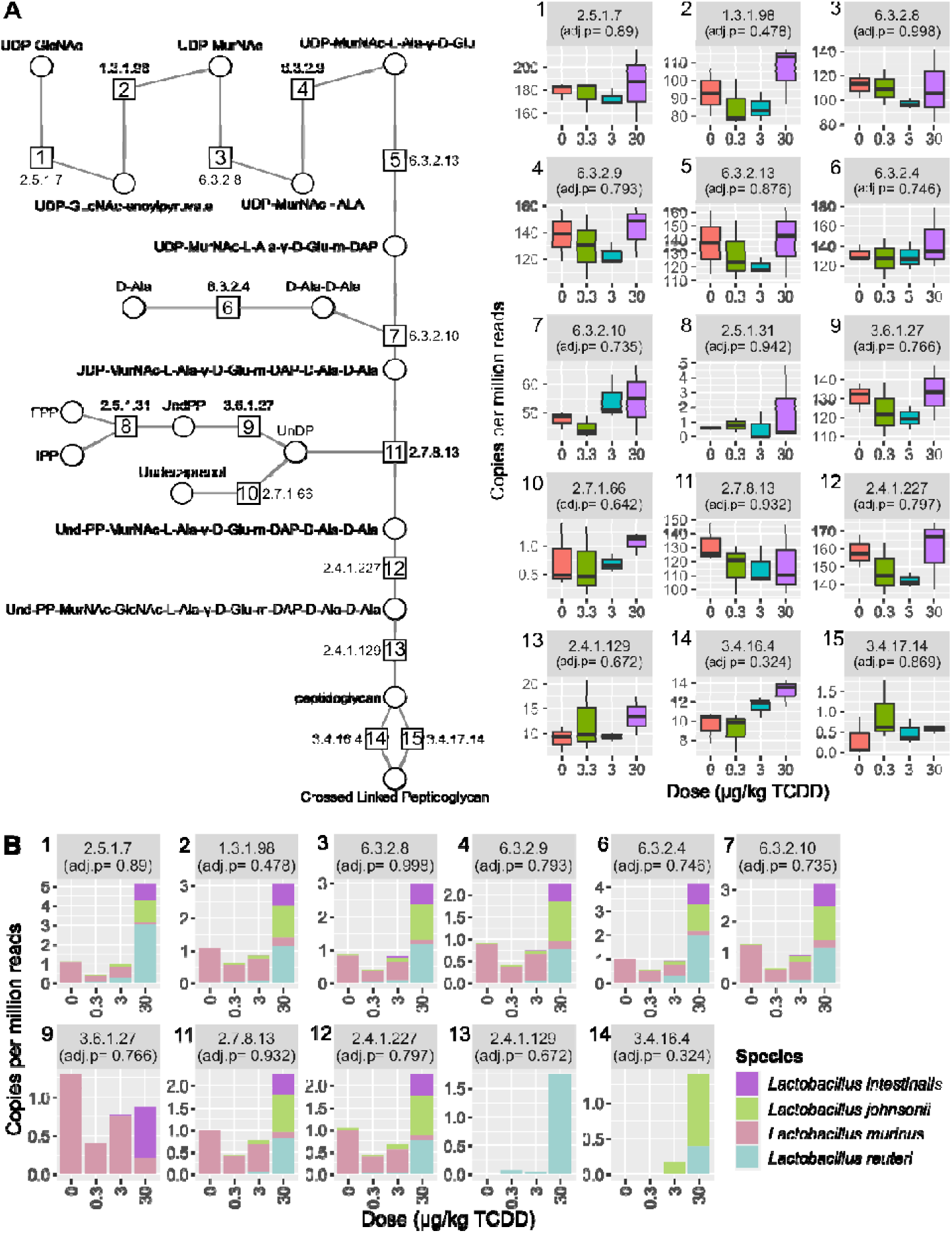
Peptidoglycan biosynthesis was unchanged by TCDD. **(A)** Relative abundance of peptidoglycan biosynthesis EC numbers identified in the metagenomic dataset. **(B)** Relative abundance of only Lactobacillus species classified to peptidoglycan biosynthesis EC numbers. Individual boxplots are numbered with the EC number matching the enzymatic step in pathway schematic. Adjusted p-values (adj.p) were determined by MAASLIN2. Abbreviations: UDP-N-acetyl-alpha-D-glucosamine (UDP-GlcNac), UDP-N-acetylmuramate (UDP-MurNAc), UDP-N-acetyl-alpha-D-muramoyl-L-alanine (UDP-MurNAc-ALA), UDP-N-acetyl-alpha-D-muramoyl-L-alanyl-D-glutamate(UDP-MurNAc-Ala-D-Glu), UDP-N-acetylmuramoyl-L-alanyl-gamma-D-glutamyl-meso-2,6-diaminopimelate (UDP-MurNAc-Ala-D-Glu-m-DAP), D-Alanyl-D-alanine (D-Ala-D-Ala), UDP-N-acetylmuramoyl-L-alanyl-D-glutamyl-6-carboxy-L-lysyl-D-alanyl-D-alanine (UDP-MurNAc-Ala-D-Glu-m-DAP-D-Ala-D-Ala), Undecaprenyl-diphospho-N-acetylmuramoyl-L-alanyl-D-glutamyl-meso-2,6-diaminopimeloyl-D-alanyl-D-alanine (Und-PP-MurNAc-Ala-D-Glu-m-DAP-D-Ala-D-Ala), Undecaprenyl-diphospho-N-acetylmuramoyl -(N-acetylglucosamine)-L-alanyl-D-glutamyl-meso-2,6-diaminopimeloyl-D-alanyl-D-alanine (Und-PP-MurNAc-GlcNAc-Ala-D-Glu-m-DAP-D-Ala-D-Ala)

However, serine-type D-Ala-D-Ala carboxypeptidase (Figure 6a, EC 3.4.16.4, step 14), responsible for peptidoglycan polymer crosslinking, trended upwards. Additionally, most peptidoglycan biosynthesis EC numbers had annotations to *L. reuteri* (Figure 6b). Overall, TCDD did not alter peptidoglycan synthesis related gene levels.

*De novo* menaquinone biosynthesis requires chorismate and the addition of a polyprenol diphosphate (i.e., geranyl-geranyl diphosphate) (Figure 7a). Two alternative pathways exist for menaquinone biosynthesis, the o-succinylbenzoate or futalosine route[45]. Only a few EC number annotations were detected for the futalosine pathway (EC 4.2.1.151 and EC 2.5.1.120), while all EC numbers were identified for the complete o succinylbenzoate menaquinone pathway (Figure 7a) In the mouse cecum dataset species contributing to o-succinylbenzoate menaquinone biosynthesis pathway included *Escherichia coli*, several Bacteroides (*e.g*., *Bacteroides vulgatus* and *Bacteroides caecimuris*), and Lactobacillus species (*e.g*., *L. reuteri*) (Figure 7b and Table S6). No one species was annotated to the entire set of enzymes needed for *de novo* biosynthesis from chorismate, however *B. vulgatus* was annotated for 6 out of 9 genes in the pathway (Table S6). O-Succinylbenzoate synthase (Figure 6a, EC 4.2.1.113, step 4) was increased by 30 µg/kg TCDD with *L. reuteri* being the major contributor to relative abundance (Figure 7, step 4). Lactobacillus species annotated to menaquinone biosynthesis included *L. reuteri, L. murinus*, and *L. johnsonii*. Among annotated menaquinone biosynthesis EC numbers, *L. reuteri* was among the identified Lactobacillus species that had the highest relative abundance and most menaquinone EC annotations (Figure 7b). *L. reuteri* also had annotations in samples for EC numbers involved in the final steps of the shikimate pathway responsible for chorismate biosynthesis (Table S7).

**Figure 7.**
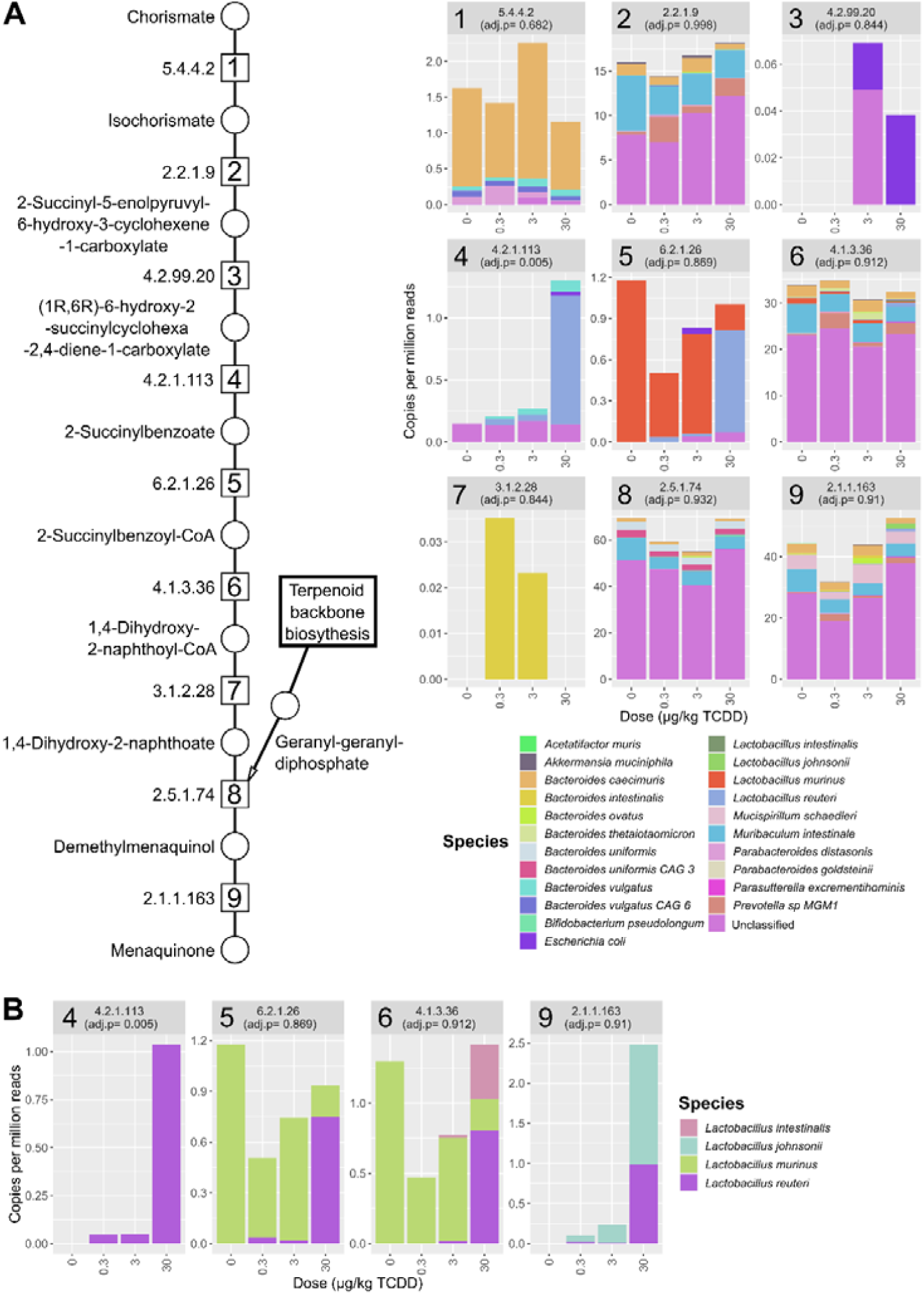
TCDD-elicited effects on menaquinone biosynthesis. **(A)** Relative abundance of menaquinone biosynthesis EC annotations identified in the metagenomic dataset. Individual stacked bar plots are labeled with the EC number matching the enzymatic step in pathway schematic. Stacked bar plots of annotated EC numbers involved in menaquinone biosynthesis. Values are mean relative abundance (n=3) classified to respective species and in cecum samples from male C57BL/6 mice following oral gavage with sesame oil vehicle or 0.3, 3, or 30 µg/kg TCDD every 4 days for 28 days. **(B)** Menaquinone biosynthesis EC numbers classified to Lactobacillus species in the cecum metagenomic datasets. Adjusted p-values (adj. p) were determined by Maaslin2 R package.

In cirrhosis samples, several EC numbers representing the initial menaquinone biosynthesis steps were also increased in compensated and decompensated patients (Figure 8, steps 1,3,4-5) including o-succinylbenzoate synthase (Figure 8, EC 4.2.1.113, step 4). However, *L. reuteri* was not among species classified to this EC number. Species classified to all EC numbers comprising the complete pathway included *E. coli*, and Klebsiella species such as *K. pneumoniae* and Citrobacter species. *L. reuteri* was not annotated to any menaquinone biosynthesis genes in healthy or compensated patients, but several EC numbers in the decompensated group (EC 6.2.1.26, 4.1.3.6, and 2.1.1.163) which are involved in later stages of menaquinone biosynthesis (Table S8).

**Figure 8.**
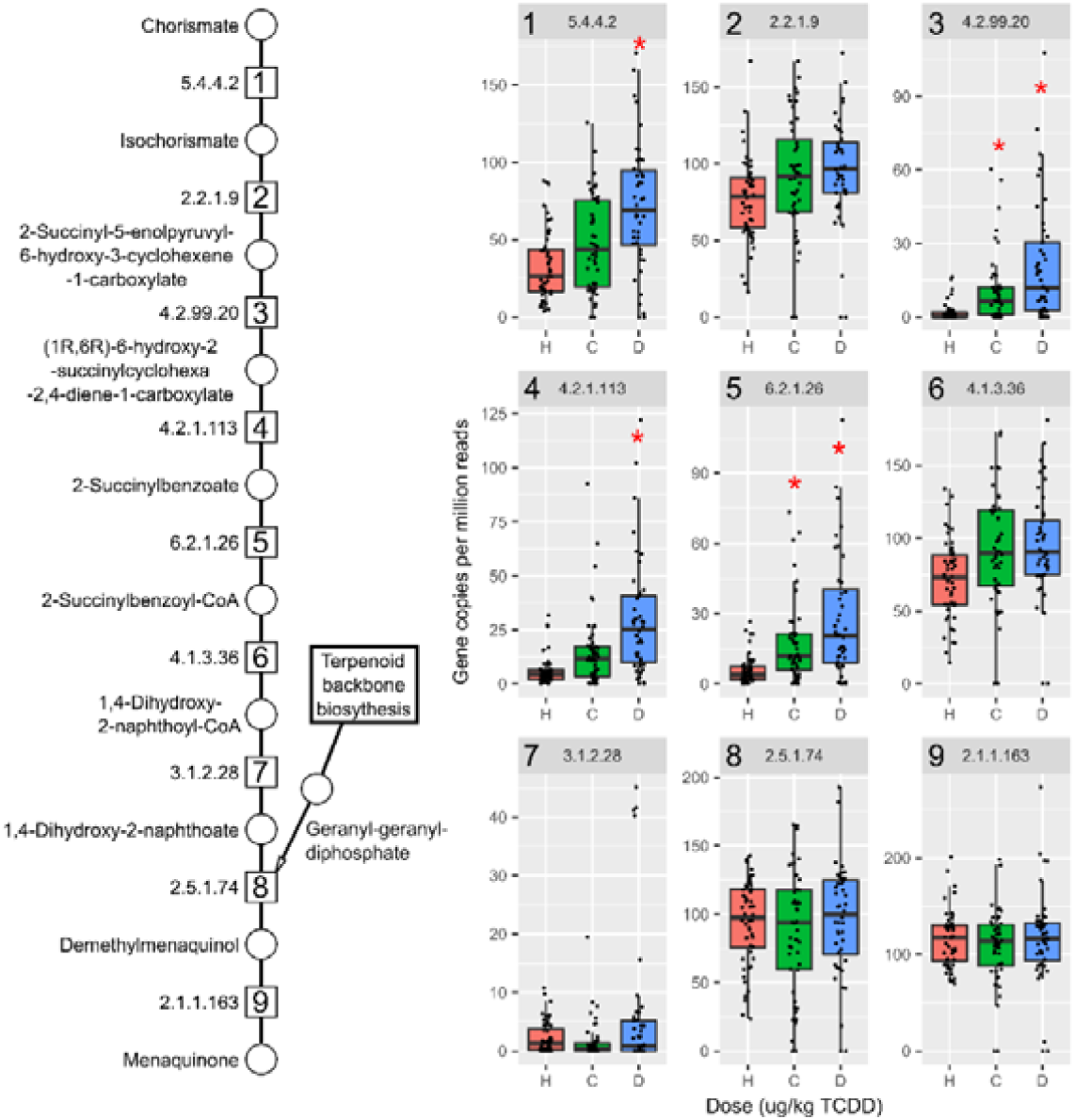
Menaquinone biosynthesis genes are increased in cirrhotic patients. Humann3 analysis of fecal metagenomic dataset of patients with healthy (H, red, n=52), compensated (C, green, n=48), or decompensated (D, blue,n=44) liver cirrhosis diagnosis for EC numbers in menaquinone biosynthesis. Individual box plots are labeled with the EC number matching the enzymatic step in pathway schematic. Significance is denoted with a red asterisk (*; adjusted p-values < 0.05) with the healthy group as reference.

## 3. Discussion

Previous studies have reported that TCDD elicited NAFLD-like pathologies, dysregulated bile acid metabolism and gut microbiome dysbiosis [9,11,15,29,31]. This study further elucidated the shifts in the gut microbiota associated with TCDD treatment using shotgun metagenomic sequencing. We show TCDD dose-dependently shifted the gut microbiota composition by enriching for Lactobacillus species, consistent with hepatic disruption of host and microbial bile acid metabolism. In addition, TCDD enriched for genes involved in mevalonate dependent isoprenoid precursor biosynthesis and menaquinone biosynthesis, crucial for microbial cell growth and survival. Over-representation of these microbial associated pathways were also identified in human cirrhosis stool metagenomics datasets.

TCDD-elicited gut dysbiosis is in agreement with observed effects in published *in vivo* studies following treatment with endogenous (*i.e*., FICZ) and exogenous (*i.e*., TCDD and TCDF) AhR agonists[8,9,11,29–31]. More specifically, we observed an increased Firmicutes/Bacteroides ratio with dose-dependent increases in Lactobacillus species[29,31]. Lactobacillus species are often associated with NAFLD, with increased abundances in patients with diabetes and liver fibrosis[46]. Probiotic Lactobacillus species including *L. reuteri* supplementation is also reported to alleviate NAFLD pathologies by reducing steatosis[47], fibrosis[48], insulin resistance[49] and serum cholesterol levels[50]. However, Lactobacillus species supplementation may also exacerbate fibrosis[51]. In humans and mice, *L. reuteri* supplementation can modulate the gut microbiota and alter bile acid metabolism. *L. reuteri* enrichment also approached comparable levels compared to samples from humans and mice administered probiotic supplementation[52,53]. We observed a species-specific increase of *L. reuteri* with a concurrent decrease in *L. murinus* suggesting shifts in Lactobacillus composition at the species and/or strain levels. Further, decreased abundance of *L. murinus* has been reported in human NAFLD[54]. Other taxa enriched following treatment included *Turicibacter sanguinis*, an anaerobic gram-positive bacillus commonly found in animals, including humans[55]. Interestingly, *T. sanguinis* has been shown to deconjugate bile acids and metabolize serotonin affecting lipid and steroid metabolism[55,56]. Quantitative trait locus analysis correlated *T. sanguinis* abundance with cholic acid levels and expression of the intestinal bile acid transporter *Slc10a2*[55]. Both cholic acid levels and *Slc10a2* expression are dose-dependently increased by TCDD[9]. Consequently, the dose-dependent taxonomic shift in Lactobacillus and Turicibacter species known to deconjugate conjugated bile acids is consistent with increased levels of secondary bile acids following TCDD treatment.

Some host relevant intestinal health and homeostatic effects can be attributed to Lactobacillus species mediated by bile salt hydrolases (*BSH*s) which are responsible for deconjugation reactions, the gateway step for conversion of conjugated primary bile acid to secondary bile acids [57]. A majority of Lactobacillus species possess *BSH*s, often containing multiple different gene copies within their genome, some with different bile acid substrate preferences[34,39]. However, the presence of *bsh* sequences does not simply infer bile acid tolerance as growth inhibition and reduced fitness is also possible depending on the conjugated or deconjugated bile acids present and/or BSH specificity.[34,39,58] For example, *L. gasseri bsh* knockout mutants exhibit increased fitness compared to wild type strains[39]. Interestingly, *L. gasseri bsh* sequences were not identified despite increased *L. gasseri* abundance following TCDD treatment. Our *bsh* analysis also found TCDD enriched Lactobacillus-associated sequences that may impart bile acid tolerance. For example, the *bsh* sequence enriched by TCDD annotated to *L. johnsonii* (RefSeq ID: EGP12391) (Table S3) exhibited higher substrate specificity for glycine over taurine conjugated bile acids[59,60]. In a companion study using the dose response and treatment regimen, Fader *et al* reported TCDD increased serum DCA levels ∼80-fold, with only a ∼ 2-fold increase in serum GDCA levels[9]. In contrast, hepatic taurolithocholic acid (TLCA) levels were increased ∼233-fold while serum lithocholic acid increased only 4-fold following TCDD treatment. Moreover, glycine conjugated bile acids including GDCA are more toxic towards Lactobacillus species than taurine conjugated bile acids[34,61,62]. Increased levels of *BSH* with a substrate preference for glycine conjugated bile acid may partially explain select Lactobacillus species enrichment. Further, both TLCA and DCA are potent FXR and GPBAR1 agonists that regulate lipid, glucose and bile acid metabolism[63,64]. Consequently, shifts in microbial secondary bile acids by Lactobacillus species may play a role in TCDD elicited gut dysbiosis impacting host regulation of energy homeostasis.

Coincident with increased levels of *bsh* was the dose-dependent increase in genes from the mevalonate-dependent isoprenoid biosynthesis, the pathway also used in mammals for cholesterol biosynthesis. The MEP pathway is the predominant isoprenoid biosynthesis pathway among gut microbiota while the mevalonate-dependent pathway is only found in select bacteria including Lactobacillus and Streptococcus species[65]. The output from either pathway is farnesyl diphosphate (FPP) and geranyl diphosphate (GPP), substrates required for polyprenol biosynthesis used in menaquinone and cell wall biosynthesis. Menaquinones are utilized by bacteria for anaerobic/aerobic respiration, providing antioxidant activity with menaquinone supplementation affecting the gut microbiome[66]. In the context of *L. reuteri*, we observed genes annotated to the shikimate pathway which is responsible for chorismate biosynthesis, a precursor for aromatic amino acids and the naphthoquinone head group of menaquinone, as well as genes involved in *de novo* menaquinone biosynthesis. While the complete biosynthesis pathway was not present in *L. reuteri*, it is consistent with other metagenomic reports of incomplete menaquinone biosynthesis pathways in gut Lactobacillus species [45]. It has been proposed that Lactobacillus species may participate in later menaquinone biosynthesis steps through the uptake of intermediates such as o-succinylbenzoate from other bacteria or dietary sources[45]. In addition, the ability to utilize menaquinones for respiration is typically not associated with Lactobacillus species. However, some lactic acid bacteria including *L. reuteri* strains demonstrate the ability to respire when menaquinone and heme are supplemented[67,68].

Metagenomic analysis also identified the mevalonate-dependent pathway enrichment in fecal samples from patients with cirrhosis. The mevalonate-dependent pathway is reported to be increased in fibrosis patients with autoimmune pathologies[69]. Isoprenoid biosynthesis pathways are also elevated in the lung microbiome of cystic fibrosis patients, with the MEP pathway enriched rather than the mevalonate route[70]. The association between fibrosis and isoprenoid biosynthesis enrichment warrants further investigation in the context of potential mechanisms contributing to bacterial fitness and/or fibrosis. Increased abundance of the mevalonate-dependent biosynthesis pathway could also be a biomarker of Lactobacillus and Streptococcus proliferation that is often associated with non-alcoholic steatohepatitis (NASH)/fibrosis[22,46]. We identified enrichment of the mevalonate-dependent pathway in both mouse and human microbiomes whereas the complete pathway was primarily annotated to Streptococcus and Lactobacillus species (Table S7). Other factors such simvastatin and proton pump inhibitors (PPI) that are commonly prescribed for NAFLD patients may also impact these microbial pathways. Simvastatin, which is primarily excreted in the feces[71], is reported to reduce bacterial growth by directly inhibiting bacterial HMG-CoA synthesis while PPIs inhibit Streptococcus species growth[72–74]. These microbiome-drug interactions highlight off target effects that should be considered when investigating novel NAFLD treatments such as new drug development and/or probiotic interventions.

In addition to increased mevalonate-dependent isoprenoid biosynthesis genes in cirrhotic patients, menaquinone biosynthesis gene abundance was also increased. This suggests taxa with the ability to produce menaquinone may have a competitive advantage when intestinal environmental conditions shift during disease progression. In cirrhosis patients, *E. coli* and *B. vulgatus* were associated with genes providing a majority of the menaquinone biosynthesis capacity. These species are also increased in human NAFLD[75]. Similar to the results in mice exposed to TCDD, *L. reuteri* was associated with several menaquinone biosynthesis genes and only detected in decompensated cirrhosis patients but lacked the complete pathway (Table S10). In cirrhosis patients, it is unclear whether *L. reuteri* is participating in menaquinone metabolism and/or benefiting from increased abundance of species, like *E. coli* that are capable of producing menaquinones.

This study was designed to account for factors affecting gut microbiota analysis including coprophagia and circadian rhythm (refs). Significant shifts in taxa were observed in Lactobacillus species. However, the small group size (n=3) following adjustment for multiple testing lacked sufficient power to confirm more subtle shifts such as the 2-fold enrichment of *Lachnospiraceae A4*, an abundant community member associated with *bsh* sequences. Samples were also collected within the same Zeitgeber period to account for possible variations in relative microbiota levels due to circadian rhythm/diurnal regulation. In fact, *L. reuteri* is one gut microbiome member demonstrating changes in relative abundance in human samples due to circadian/diurnal regulation[76]. TCDD disrupted diurnal regulation of hepatic gene expression including bile acid biosynthesis genes which may contribute to *L. reuteri* enrichment[77].

Although the consequences of TCDD-elicited immune system effects on the gut microbiome were not assessed in this study, it is most likely a factor impacting *L. reuteri* enrichment. TCDD causes macrophage and dendritic cell migration out of the lamina propria with increased accumulation in the liver, possibly exacerbating hepatic inflammation and affecting intestinal immune responses[14]. The ability of *L. reuteri* to produce AhR ligands, upregulate IL-22, and associate with the mucosa and Peyer’s patches provides geographical proximity for immune/microbiome crosstalk mediated by the AhR [27,78,79]. Besides immune cell regulation, TCDD increased bone formation and decreased bone marrow adiposity[80]. Interestingly, *L. reuteri* supplementation also increased bone density, but only when mice are induced towards an inflammatory state[81]. Overall, the dose-dependent increase in *L. reuteri* levels are consistent with increased bile acid levels, disruption of circadian/diurnal regulation and increased bone density[9,77,80,81].

In summary, Lactobacillus species were dose-dependently increased following AhR activation by TCDD, concurrent with the increase in *bsh* genes and increased primary and secondary bile acids. Specifically, *L. reuteri*, a keystone gut microbiome species is involved in microbial metabolism of bile acids and AhR ligands. The large and uniform enrichment of *L. reuteri* in this study also suggests environmental pressures such as increased levels of bile acids and antimicrobial peptides elicited by AhR activation may provide a complementary niche for *L. reuteri* that possess a gene repertoire not found in the closely related *L. murinus*. We also provide evidence on how *L. reuteri* metabolism could impact AhR, FXR and GPBAR1 signaling pathways, placing *L. reuteri* at the crossroads of bacterial/host interactions affecting glucose, bile acid, and immune regulation. Whether these microbial shifts in metabolism are adaptive and limit the intensity of adverse consequences or exacerbates steatosis to steatohepatitis with fibrosis progression warrants further investigation.

## 4. Materials and Methods

### 4.1 Animal treatment

Postnatal day 25 (PND25) male C57BL/6 mice weighing within 10% of each other were obtained from Charles River Laboratories (Kingston, NY) and housed and treated as previously described[9]. Briefly, mice were housed in Innovive Innocages (San Diego, CA) containing ALPHA-dri bedding (Shepherd Specialty Papers, Chicago, IL) in a 23°C environment with 30–40% humidity and a 12 hr/12 hr light/dark cycle. Aquavive water (Innovive) and Harlan Teklad 22/5 Rodent Diet 8940 (Madison, WI) were provided *ad libitum*. The rodent diet is a fixed formula complete diet with an energy density of 3.0 kcal/g and a nutrient ingredient composition including 22% protein, 5.5% fat, and 40.6% carbohydrate. Mice (PND29) were orally gavaged at the beginning of the light cycle (between Zeitgeber time 0-3) with 0.1 ml sesame oil vehicle (Sigma-Aldrich, St. Louis, MO) or 0.3, 3 and 30 μg/kg body weight TCDD (AccuStandard, New Haven, CT) every 4 days for 7 total exposures (n=3 per treatment group). The study was conducted in three cohorts with mice housed separately among treatment groups for a total of 9 mice per treatment group. In each cohort, three mice were housed per treatment group and one mouse was randomly selected from each treatment group per cohort (n=3 per treatment group for the metagenomic analysis) to account for coprophagia and ensure reproducibility. The first gavage was administered on day 0 of the study. On day 28, vehicle- and TCDD-treated mice (fasted for 6 hr with access to water) were weighed and euthanized between Zeitgeber time 0-3. Upon collection, cecums were immediately flash frozen in liquid nitrogen and stored at -80°C until analysis. All animal handling procedures were performed with the approval of the Michigan State University (MSU) Institutional Animal Care and Use Committee.

### 4.2 Metagenomic sequencing

Microbial DNA from cecum contents (∼25 mg) was extracted using the FastDNA spin kit for soil (SKU 116560200, MP Biomedicals, Santa Ana, CA). Extracted DNA was submitted to Novogene (Sacramento, CA) for quality control, library preparation, and 150-bp paired-end sequencing at a depth 136-157 million reads using an Illumina NovaSeq 6000. Reads aligning to the C57BL/6 *Mus musculus* genome (NCBI genome assembly: GRCm38.p6) were identified, flagged, and removed using bowtie2[82], SamTools[83] and bedtools[84]. For human metagenomic analysis, reads were filtered against the human genome (NCBI genome assembly: GRCh37/hg19) using the Kneaddata bioinformatics tool developed at the Huttenhower Lab (https://github.com/biobakery/kneaddata).

### 4.3 Metagenomic taxonomic analysis

Kaiju was used for taxonomic analysis of mouse cecum metagenomic dataset. The reference database used was the progenomes database downloaded from the kaiju webserver (https://kaiju.binf.ku.dk/database/kaiju_db_progenomes_2020-05-25.tgz). Multivariate association between dose and taxonomy relative abundances used Maaslin2 (https://github.com/biobakery/Maaslin2)[85] with the following default settings used: normalization (total sum scaling), analysis method (general linear model), and Benjamini-Hochberg multiple test correction. Adjusted p-values for Maaslin2 analysis used dose (sesame oil vehicle (0), 0.3, 3, or 30 μg/kg TCDD) as the fixed effect which was treated as continuous variable and the vehicle set for reference. For comparison of taxonomy between vehicle and 30 μg/kg TCDD treatment groups, DeSeq2 was used to determine adjusted p-values using default settings[86].

### 4.4 Metagenomic functional analysis

The HUMAnN 3.0 bioinformatic pipeline[87] was used with default settings to classify reads to UniRef90 protein identifications using UniProt’s UniRef90 protein data base (January, 2019). Reads aligned to UniRef90 identifications were mapped to enzyme commission (EC) number entries using the humann_regroup_table tool. Read abundance was normalized to gene copies per million reads (CPM) using the human_renorm_table tool. Multivariate association between dose and enzyme commission number relative abundance used Maaslin2 with same settings used for taxonomy analysis.

Xander (a gene-targeted assembler, https://github.com/rdpstaff/Xander_assembler) was used to annotate and quantify bile salt hydrolase sequences with the following settings: k-mer size=45, filter size=35, minimum assembled contig bit score=50, and minimum assembled protein contigs=100 [88]. Reference DNA and protein *bsh* sequences used for Xander were downloaded from FunGenes Gene Repository and are listed in supplementary material (Table S9 and Table S10)[89]. For RefSeq *bsh* sequences analysis, relative abundance was determined by normalizing to total abundance of *rplB* sequences also determined by Xander per sample. Significance was determined with Maaslin2 with same settings used for taxonomy analysis

Human metagenomic data from the European Bioinformatics Institute European Nucleotide Archive under accession number PRJEB6337 (https://www.ebi.ac.uk/ena/data/view/PRJEB6337) was analyzed using the same HUMAnN 3.0 pipeline as cecum metagenomic data. Fecal shotgun metagenomic samples from Chinese patients were defined as healthy (n=52) or cirrhotic with subclassifications of compensated (n=48) or decompensated (n=44) by the authors[40]. Cirrhosis was diagnosed by either biopsy, clinical evidence of decompensation or other metrics including radiological evidence of liver nodularity and intra-abdominal varices in a patient with chronic liver disease[40]. The subclassification was used as fixed effect for analysis with healthy as the reference category. Again, Maaslin2 was used with settings used for functional analysis with diagnosis as a fixed effect with healthy diagnosis as reference to determine adjusted p-values for compensated and decompensated patient designations.

## Supplementary Materials

The following are available online at www.mdpi.com/xxx/s1, Figure S1: Stacked bar plots of annotated EC number mean relative abundance and classified species involved in isoprenoid biosynthesis in cecum of TCDD-exposed mice, Figure S2: Stacked bar plots of mean relative abundance of annotated EC numbers and classified to Lactobacillus species involved in isoprenoid biosynthesis in cecums of TCDD-exposed mice, Figure S3: Stacked bar plots of annotated EC number mean relative abundance classified to respective species and involved in mevalonate dependent isoprenoid biosynthesis in cirrhotic patients, Table S1: Relative abundance of significantly changed *bsh* sequences in mouse cecums, Table S2: Significant Uniref90 annotations in murine cecum metagenomic dataset, Table S3: Enzyme commission numbers (EC) that were significant and annotated to at least *L. reuteri* in cecums of TCDD-exposed mice, Table S4: Enzyme commission numbers (EC) with species annotated to mevalonate-dependent IPP pathway for cirrhotic patients, Table S5: Total number of unique EC numbers determined to have greater than zero relative abundance in respective pathway (mevalonate and MEP isoprenoid biosynthesis, and menaquinone biosynthesis) in human cirrhosis patients, Table S6:Total number of unique EC numbers determined to have greater than zero relative abundance in respective pathway (mevalonate and MEP isoprenoid biosynthesis, and menaquinone biosynthesis) in TCDD-exposed mice, Table S7: Mean relative abundance denoted in counts per million (CPM) of EC annotations that were classified to *L. reuteri* and associating with chorismate biosynthesis., Table S8: EC numbers with relative abundance of species to menaquinone biosynthesis pathway for cirrhotic patients, Table S9: Nucleotide sequences used as reference in Xander *bsh* analysis, Table S10. Protein sequences used as reference in Xander *bsh* analysis.

## Author Contributions

Conceptualization, R.F. and T.Z.; methodology, R.F, T.Z.; writing—original draft preparation, R.F..; writing—review and editing, R.F, T.Z.; funding acquisition, T.Z; formal analysis, R.F.; visualization, R.F..; data curation, R.F; Methodology R.F, T.Z; validation, R.F.; project administration, T.Z; investigation, R.F., T.Z; All authors have read and agreed to the published version of the manuscript.

## Institutional Review Board Statement

All animal handling procedures were performed with the approval of the Michigan State University (MSU) Institutional Animal Care and Use Committee, in accordance with ethical guidelines and regulations under IACUC ID: PROTO201800043.

## Informed Consent Statement

Informed consent was obtained from all subjects involved in the human metagenomics study[40].

## Data Availability Statement

Quality filtered mouse metagenomic data from this study can be found at the Sequence Read Archive (SRA) (https://www.ncbi.nlm.nih.gov/sra) under the accession ID PRJNA719224. The human fecal metagenomic data presented in this study is openly available and can be found at the SRA under the accession number PRJEB6337[40].

