## Supplementary Figures for "Aryl hydrocarbon receptor (AhR) activation by 2,3,7,8-tetrachlorodibenzo-*p*-dioxin (TCDD) dose-dependently shifts the gut microbiome consistent with the progression of steatosis to steatohepatitis with fibrosis"

**Supplemental Material**

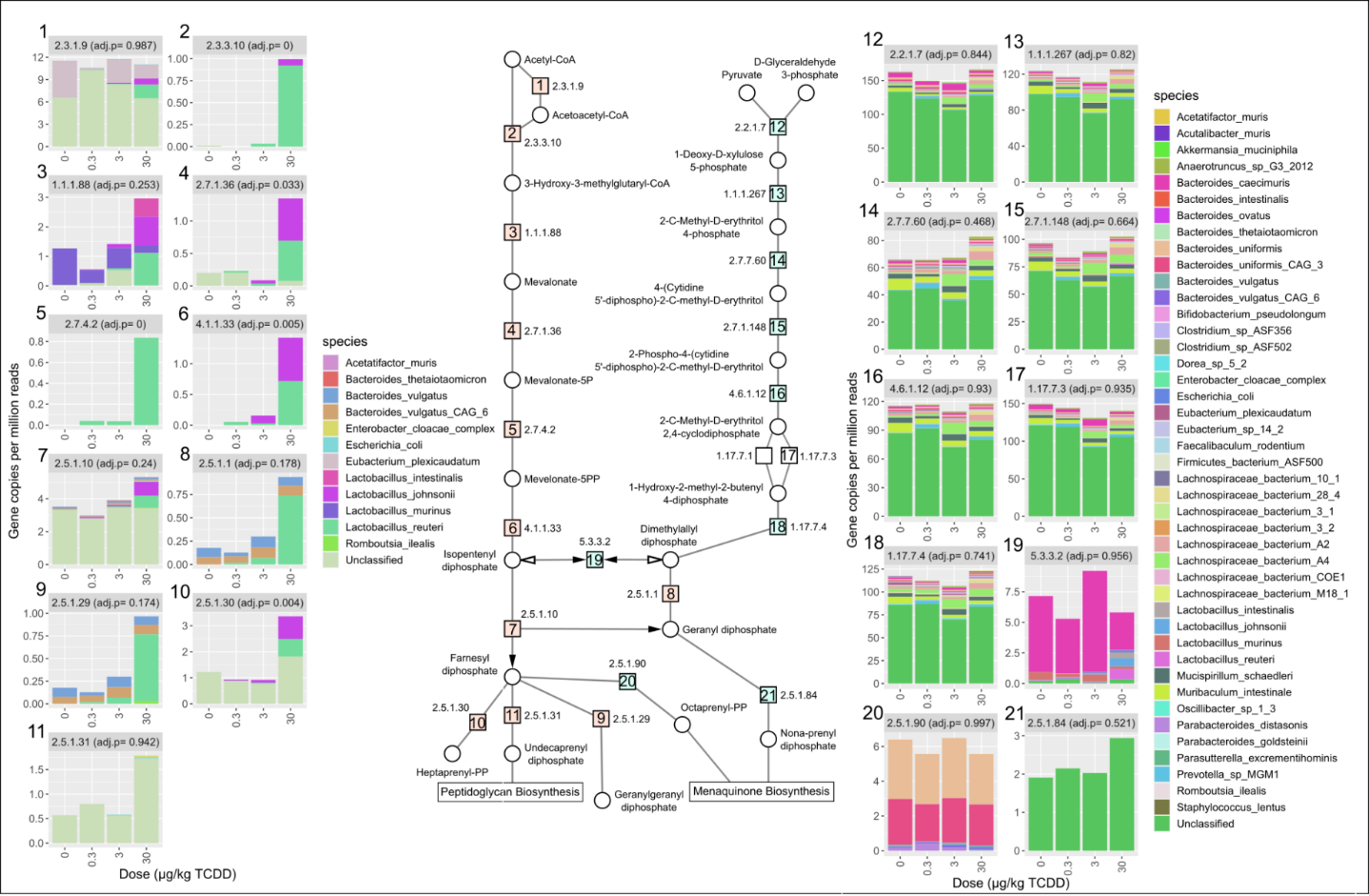

**Figure S1.** Stacked bar plots of annotated EC number mean relative abundance and classified species involved in isoprenoid biosynthesis in cecum of TCDD-exposed mice. Male C57BL6 mice were orally gavaged with sesame oil vehicle or 0.3, 3, or 30 µg/kg TCDD every 4 days for 28 days (n=3). Numbers beside each bar plot correlate with numbered EC number provided in the center schematic of isoprenoid biosynthesis. Adjusted *p*-values (adj. p) were determined by the Maaslin2 R package.

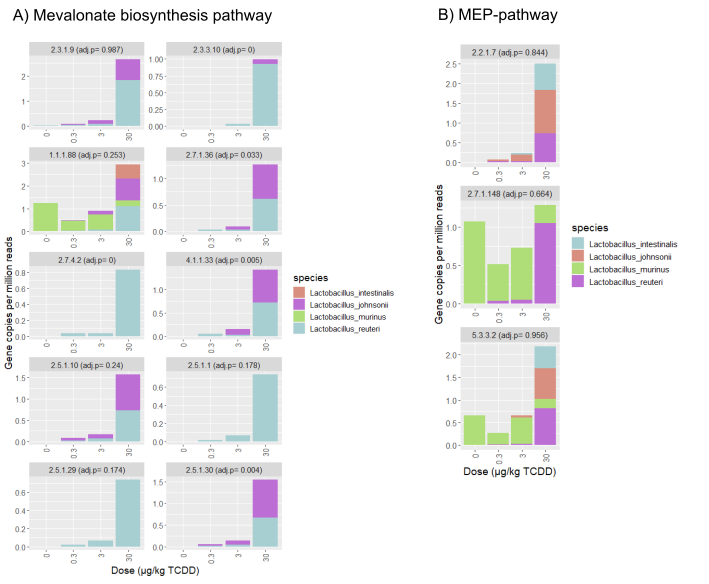

**Figure S2.** Stacked bar plots of mean relative abundance of annotated EC numbers and classified to Lactobacillus species involved in isoprenoid biosynthesis in cecum of TCDD-exposed mice. Male C57BL6 mice were orally gavaged with sesame oil vehicle or 0.3, 3, or 30 µg/kg TCDD every 4 days for 28 days (n=3). Adjusted *p*-values (adj. p) were determined by the Maaslin2 R package and denote significance.

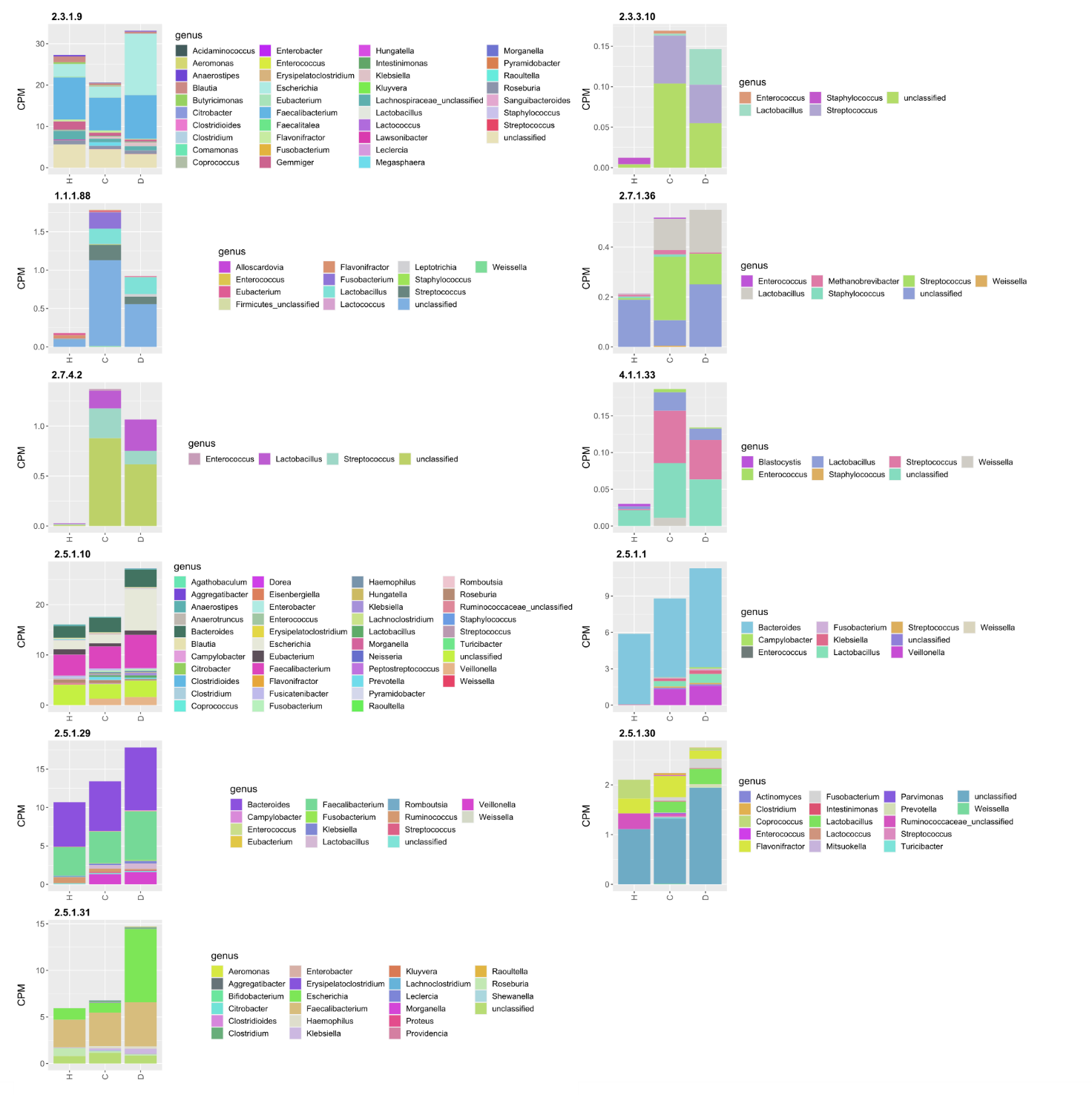

**Figure S3.** Stacked bar plots of annotated EC number mean relative abundance classified to respective genus and involved in mevalonate-dependent isoprenoid biosynthesis in cirrhosis patients. Fecal metagenomic samples were from healthy (H, n=52), compensated (C, n=48) or decompensated (D, n=44) diagnostically defined groups.
